# Histone mark age of human tissues and cells

**DOI:** 10.1101/2023.08.21.554165

**Authors:** Lucas Paulo de Lima Camillo, Muhammad Haider Asif, Steve Horvath, Erica Larschan, Ritambhara Singh

**Affiliations:** School of Biological Sciences, University of Cambridge, UK; School of Clinical Medicine, University of Cambridge, UK; Department of Computer Science, Brown University, USA; Altos Labs, Cambridge, UK; Center for Computational Molecular Biology, Brown University, USA; Department of Molecular Biology, Cell Biology and Biochemistry, Brown University, USA

**Keywords:** aging, prediction, histone modifications

## Abstract

**Background:** Aging involves intricate epigenetic changes, with histone modifications playing a pivotal role in dynamically regulating gene expression. Our research comprehensively analyzes seven key histone modifications across various tissues to understand their behavior during human aging and formulate age prediction models.

**Results:** These histone-centric prediction models exhibit remarkable accuracy and resilience against experimental and artificial noise. They showcase comparable efficacy when compared with DNA methylation age predictors through simulation experiments. Intriguingly, our gene set enrichment analysis pinpoints vital developmental pathways crucial for age prediction. Unlike in DNA methylation age predictors, genes previously recognized in animal studies as integral to aging are amongst the most important features of our models. We also introduce a pan-histone-mark, pan-tissue age predictor that operates across multiple tissues and histone marks, reinforcing that age-related epigenetic markers are not restricted to particular histone modifications.

**Conclusion:** Our findings underscore the potential of histone marks in crafting robust age predictors and shed light on the intricate tapestry of epigenetic alterations in aging.

## Background

Aging is marked by noticeable changes mainly at cellular and organismal levels, encompassing phenomena like epigenetic disturbances, genomic instability, proteostasis loss, nutrient-sensing deregulation, and dysbiosis [1, 2]. This understanding has spawned a variety of omics age predictors in fields such as epigenetics, transcriptomics, proteomics, metabolomics, and microbiotics [3, 4]. Most studies focus on on blood chemistry, transcriptomics, and DNA methylation, revealing several aging biomarkers, including those based on DNA methylation, telomere length, and proteomics [3, 5]. Blood tests, facilitated by deep neural networks, offer notable accuracy with a median absolute error of about five to six years [6, 7]. RNA sequencing contributes rich transcriptomic data for age predictors, which tend to be cell type or tissue-specific [8–12]. Cytosine methylation have emerged as the most favored molecular measurement for crafting age predictors that apply to all human tissues, with pantissue predictors achieving a median absolute error nearing four years [13–16]. Single-cell pan-tissue predictors utilizing DNA methylation have also been realized [17]. Newly, pan-mammalian epigenetic clocks applicable to all mammalian species have been presented [18]. Reflecting upon the achievements of age predictors centered on cytosine methylation, it beckons whether other epigenetic shifts, notably those anchored in histone levels, could engender mammalian aging clocks of similar precision. Clocks informed by histone marks carry potential relevance, resonating well with the histone code. Despite studies delineating the nuanced relationship between aging and histone marks [19, 20], a multifaceted mammalian age predictor rooted in histone mark data remains to be formulated.

To bridge this gap, we harness ENCODE data [21, 22] to analyze seven histone marks in human tissues and cells. We pinpoint a discernible shift from heterochromatin-linked modifications to those linked with euchromatin, corroborated by other studies [19, 23]. Furthermore, the variance in these histone modifications across genes escalates with age, hinting at an epigenetic regulatory decline. We introduce the first age predictors grounded in histone modification ChIP-Seq data. Impressively, their performance rivals that of DNA methylation age predictors, adjusted for training sample size. Our explorations divulge pivotal pathways and genes for age prediction. Developmental pathways and micro RNAs conspicuously dominate most histone modification age predictors. We also unearth that histone modifications can be broadly categorized as either activating or repressive for age predictor construction, irrespective of their unique roles. Interestingly, specific genes manifest consistent age-related trends across both these categories. Capitalizing on these insights, we present the inaugural pan-histone mark, pan-tissue age predictor.

To encapsulate, our research underscores the potential of histone modification data in age prediction. We emphasize the pivotal role of epigenetic regulation in aging and spotlight the prospective utility of histone modifications as aging biomarkers. Concurrently, we illuminate key pathways and genes pivotal to epigenetic aging.

## Results

### Dynamics of histone mark during aging

In this study, we explore publicly available chromatin-immunoprecipitation sequencing (ChIP-Seq) human data from the ENCODE project [21, 22] (Figure 1). We focused on seven key histone modifications. Three, H3K4me3, H3K27ac, and H3K9ac, are broadly associated with euchromatin; another two, H3K9me3 and H3K27me3, are broadly associated with heterochromatin; one, H3K36me3, is associated with transcription elongation and heterochromatin; and one, H3K4me1, is associated with enhancers [24, 25].

**Figure 1:**
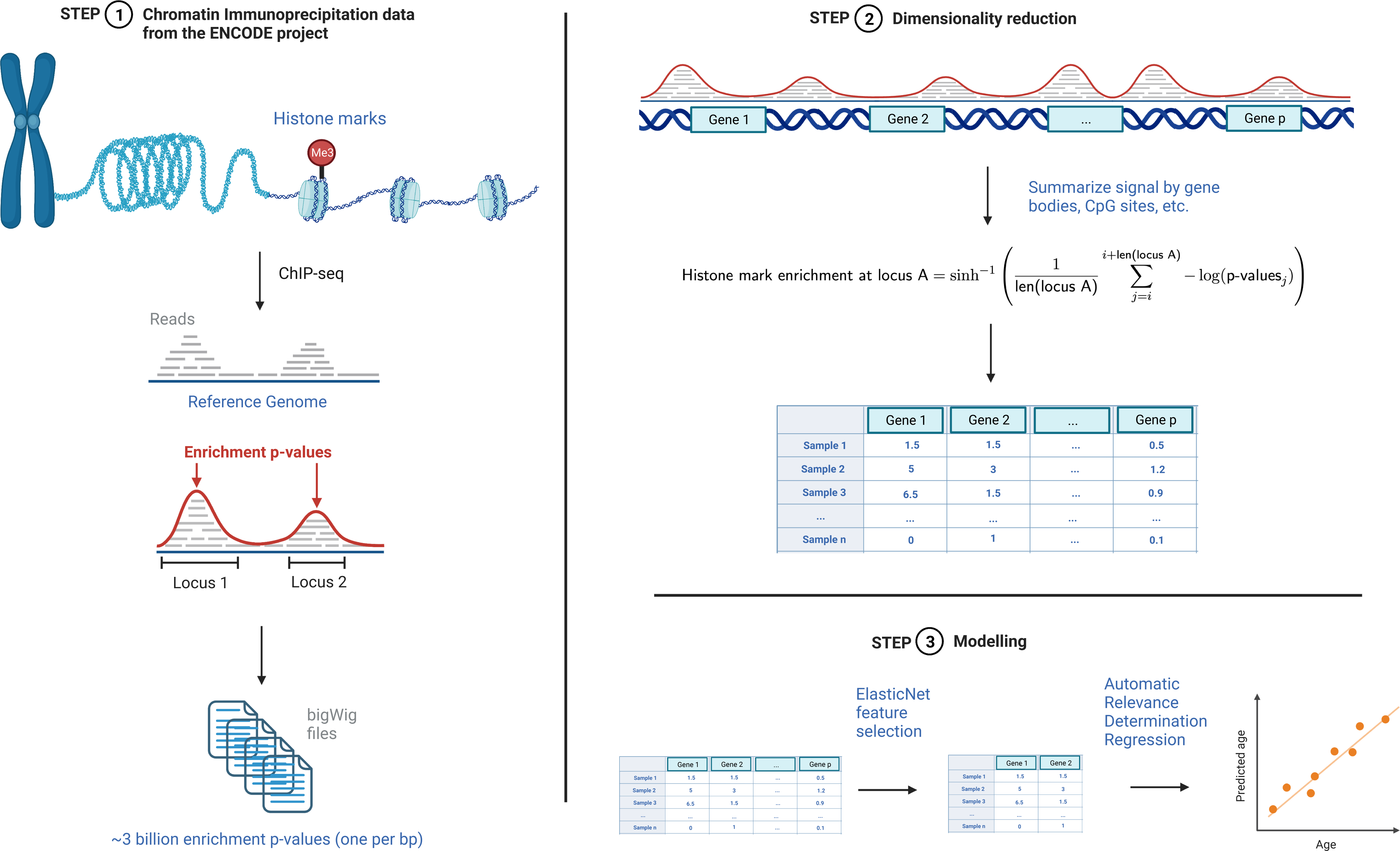
Schematic showing the main steps of creating the histone mark age predictors. First, bigWig files containing the enrichment p-values per nucleotide of chromatin immunoprecipitation samples were gathered from the ENCODE project. Then, the dimensionality of the data is reduced by summarizing the p-values into bins that represent the signal. Then, modeling is done through the application of an ElasticNet for feature selection followed by Automatic Relevance Determination regression to predict age. Image created with BioRender.

Before any attempt to create age predictors based on histone modifications, we first analyzed data derived from human tissue from the ENCODE project to understand age-related dynamics [21, 22]. We obtained a total of 1814 samples (n) from ChIP-Seq data for H3K4me3 (n = 359), H3K27ac (n = 359), H3K27me3 (n = 291), H3K4me1 (n = 264), H3K36me3 (n = 257), H3K9me3 (n = 248), and H3K9ac (n = 36). The samples represent 82 tissues with ages ranging from embryonic to 90-plus years (Supplementary Figure 2a) with a roughly equal number of males and females (48.2% males, 50.6% females, 1.2% not available). Cancerous tissue constituted 0.9% of observations (n = 17). The sequencing was performed with seven different Illumina instruments at four labs in universities across the United States. A summary of the relevant statistics can be found in Supplementary Figure 2 and attest to the breadth and diversity of the data collected.

We used the processed ChIP-Seq data files that display the probability that a genomic region is enriched for a specific histone mark compared to the control of sequencing DNA without the immunoprecipitation step. Effectively, each sample has a p-value for each single nucleotide in the genome. The lower the p-value, the higher the confidence that the locus contains the histone mark of interest. Given the high dimensionality of the data (high number of features *p* compared to the number of observations *n*), with 3 billion nucleotides in the human genome, we decided to reduce the number of features by summarizing the values across genomic regions. We opted for averaging over the gene bodies of protein-coding and noncoding genes to facilitate interpretation unless stated otherwise (minimum, median, and maximum bin sizes of 7, 3897, and 2473538 bps respectively). The Homo Sapiens Ensembl annotation 105 provided the genomic locations [26]. To summarize the values in each gene as a single feature, we averaged the negative log10 of p-values and then arcsinh-transformed to stabilize the variance (see Methods). In the end, the number of features was reduced by nearly 50 thousand, from roughly 3 billion to 62241 per sample.

We plotted our data using uniform manifold approximation and projection (UMAP) to determine whether age was a major differentiating factor in our samples. As expected, each histone modification is generally separated from the others (Figure 2a). Interestingly, however, there are two main clusters, one with the activating marks H3K4me3, H3K27ac, and H3K9ac, and another with the repressive modifications H3K9me3, H3K27me3, and the elongation mark H3K36me3. H3K4me1, typically enriched in enhancers and neither clearly activating nor repressive, is located between the two clusters. Next, we colored the UMAP plot with age rather than histone marks (Figure 2b). While there is no apparent separation between young and middle-aged samples, old samples (*>*70 years) are relatively separated from the rest of the data. We confirmed this observation by replotting UMAP stratified by modification (Figure 2c). While it is much easier to differentiate a sample based on the type of histone mark, age is relevant enough — at least in old age — to contribute towards the UMAP projection. We also plotted the data with principal component analysis (PCA) to rule out any potential artifacts from UMAP, yielding similar results (Supplementary Figures 2d-f).

**Figure 2:**
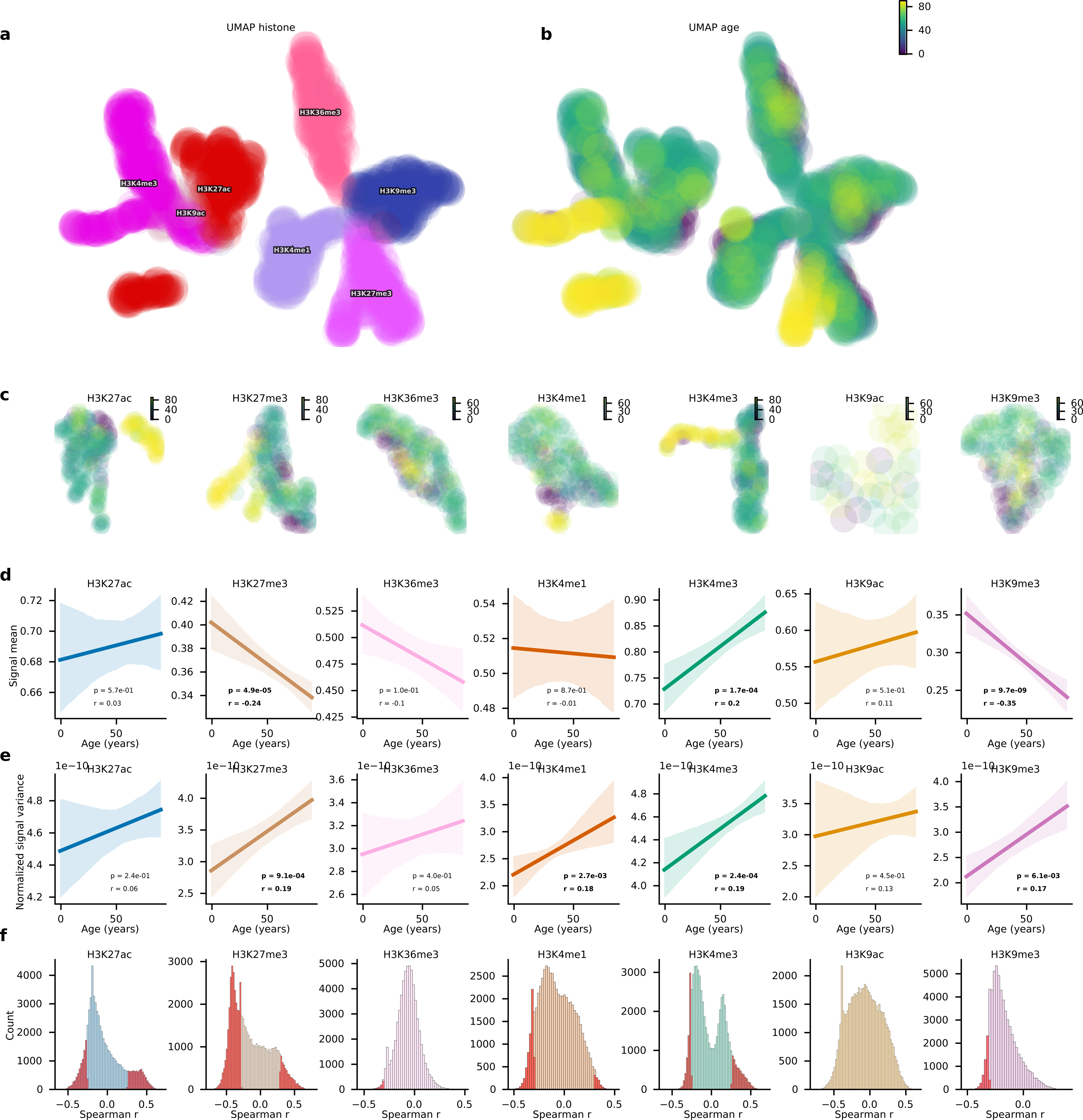
Age-related changes in seven histone modifications across human tissues. (a, b) Uniform manifold approximation and projection (UMAP) of the dataset containing the average signal of 62241 genes for 1814 samples grouped by histone modification (a) and age (b). (c) UMAP colored by age for each histone modification. (d, e) Linear regression plot with 95% confidence interval based on 1000 bootstraps for the signal mean (d) and signal variance (e) normalized by signal mean over age for each histone modification. (f) Histogram of Spearman’s correlation for each 62241 features across age per histone modification. Bins shaded in red represent statistically significant correlations (p *<* 0.05 with Bonferroni’s correction).

After broadly analyzing the data through low-dimensional projections, we focused on uncovering age-related trends. It has been widely reported that aging is accompanied by loss of heterochromatin and activation of constitutively repressed genes [19, 23]. Indeed, the mean signal of all three repressive histone modifications has a negative Pearson’s correlation (r) with age (Figure 2d), with H3K9me3 (Pearson’s r = -0.35, p-value = 9.7e-9) and H3K27me3 (Pearson’s r = -0.24, p-value = 4.9e-5) reaching significance. Likewise, the mean signal of all three activating histone marks has a positive Pearson’s correlation with age, with H3K4me3 (Pearson’s r = 0.2, p-value = 1.7e-4) reaching significance. The mean signal of H3K4me1 barely changes with age, with Pearson’s r of only -0.01. Our results add to the evidence from several previous studies indicating loss of heterochromatin with aging.

Another interesting metric to track across aging is how variable the histone modification signal becomes, as previous studies have reported an increase in entropy during aging in DNA methylation [27–29]. While the entropy calculation for an unbound histone mark enrichment is not as straightforward as for DNA methylation, the signal variance normalized by its mean can give insights into the increased variability during aging. Interestingly, Pearson’s correlation is indeed positive for all seven histone marks (Figure 2e), with H3K4me3 (Pearson’s r = 0.19, p-value = 2.4e-4), H3K4me1 (Pearson’s r = 0.18, p-value = 2.7e-3), H3K9me3 (Pearson’s r = 0.17, p-value = 6.1e-3), and H3K27me3 (Pearson’s r = 0.19, p-value = 9.1e-4) reaching significance. The broad increase in normalized signal variance with age suggests that any tight regulation to maintain histone marks to specific genomic regions becomes less effective with spillover to other loci.

While the data so far point towards robust age-related trends, we wanted to determine whether the genes’ signals correlate with age. For each of the 62241 genes, we calculated the correlation coefficient with respect to age and plotted it on a histogram (Figure 2f). We chose Spearman’s r over Pearson’s r given that multiple age-related changes are non-linear, so a non-parametric coefficient is best suited to detect such correlations. As shown by the red shade, a surprisingly large proportion of genes significantly correlate with age (p-value *<* 8.0×10-7, i.e., p-value *<* 0.05 with Bonferroni correction for the 62241 genes). As expected for the repressive histone marks, many more genes have a negative rather than positive coefficient. Surprisingly, however, the same is true for the three activating histone marks. Overall, given a large number of genes whose histone marks significantly correlated with age, it was likely that we could develop a histone mark age predictor from our data.

### Performance of pan-tissue histone mark age predictors

Given the dynamics of histone modifications we observed during aging, we set out to create age predictors. Given the relatively small number of samples, we opted for a 10-fold nested cross-validation setup. Nine folds are used for internal 9-fold cross-validation to select the appropriate hyperparameters. Then, a predictor is trained on these nine folds with the best hyperparameter and is tested in the remaining external fold. This process is repeated ten times. It is worth emphasizing that samples originating from the same biosample were not split into different folds, as this may have artificially inflated the performance (Supplementary Figure 1).

To create an apt age predictor, we had three requirements: (1) the approach can suitably handle data with high dimensionality (*p* features *>> n* samples), (2) the approach is robust to technical variation and experimental noise, and (3) the approach is easily interpretable. We introduced a feature-reduction step to fulfill the first requirement by training an ElasticNet model and selecting features with non-zero coefficients (*p^′^*) [30]. This is the only step in the overall age predictor with a hyperparameter (*λ*), representing the strength of regularization. Moreover, having an initial set of reduced features allow us to easily interpret the most important genes and pathways through gene set enrichment analysis. For the second requirement, it has been shown that PCA can vastly improve the reliability of DNA methylation epigenetic age predictors by removing technical noise [31]. Therefore, we transformed the data using PCA calculated with a truncated support vector decomposition, generating (*p^′^ −* 1) principal components. Finally, we used an automatic relevance determination regression (ARD), a form of regularized Bayesian regression, for the last requirement. It can easily be interpreted as, similarly to linear regression, each feature has a coefficient representing the model’s weight. In addition, it provides an uncertainty value for each prediction. Choosing a different model to ElasticNet after feature selection and noise reduction also avoids the issue of double dipping. While we and others have previously shown that deep learning can improve the performance and interpretation of pan-tissue DNA methylation epigenetic age predictors [14, 32], we opted for the aforementioned machine learning approach given the low number of samples. For more details of our modeling approach, see Methods.

Since chromatin immunoprecipitation is noisy and highly dependent upon the quality of the antibody [33], we expected a good but not impressive performance. We measured the performance of the age predictors using the following metrics: Pearson’s correlation coefficient (r), median absolute error (MAE), and root-mean-square error (RMSE). Surprisingly, all performed exceedingly well (Figure 3a), except for the H3K9ac age predictor -likely given the small sample size of 36. The H3K4me3 age predictor, in particular, was the best performer, with r=0.94, MAE= 4.31, and RMSE=8.74. Though the setup is not directly comparable, it is remarkable that this pan-tissue histone mark age predictor has similar reported performance compared to some of the most used DNA methylation age predictors [13, 27]. In summary, histone mark age predictors have remarkably low prediction errors.

**Figure 3:**
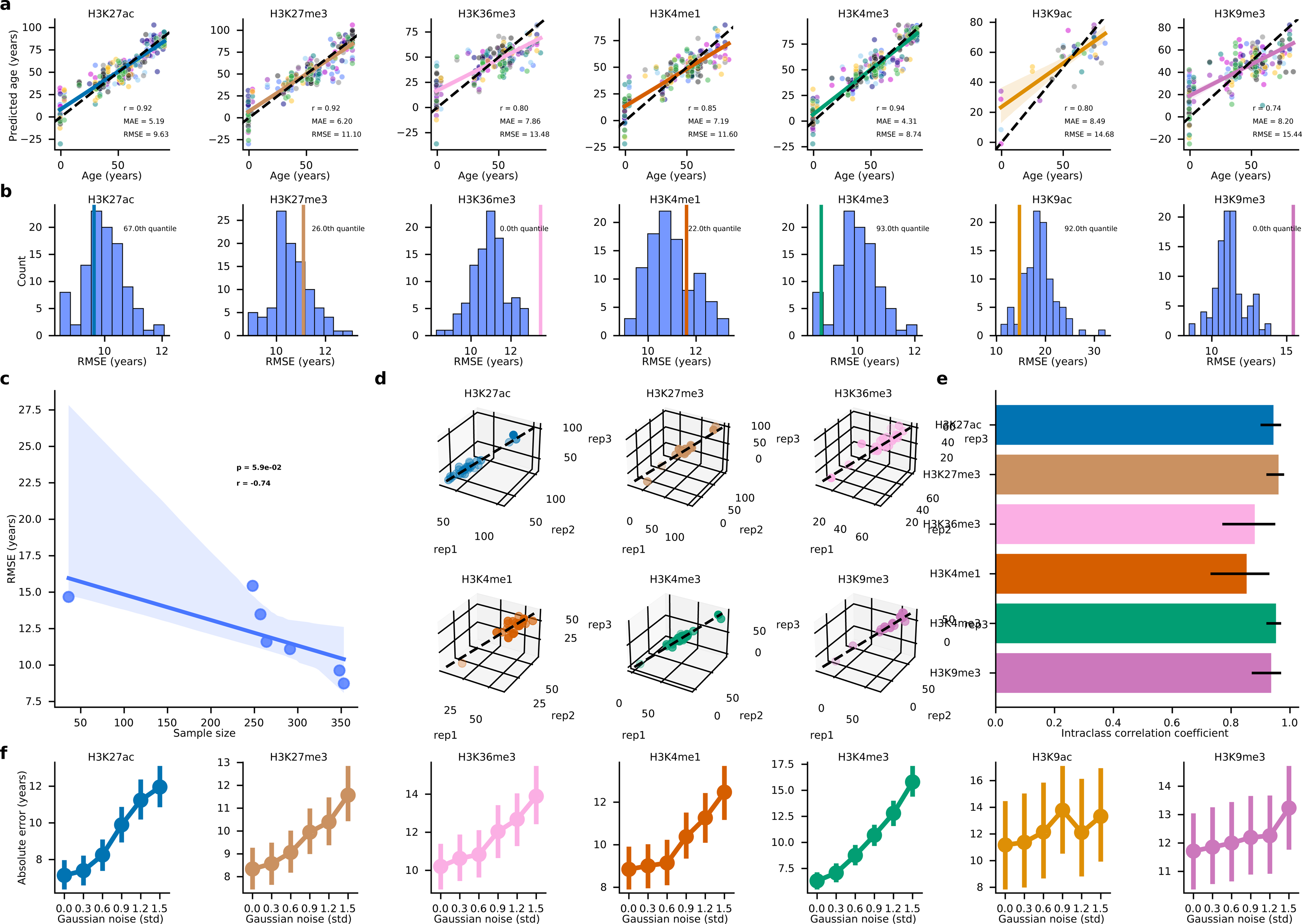
Performance of pan-tissue histone mark age predictors. (a) Scatter plot of the predicted age versus real age of each histone mark age predictor using genes as features. Each of the 10 test folds of the nested cross-validation is shown in a different color. A dotted black line representing x=y is shown alongside a colored, solid regression line with its 95% confidence interval based on 1000 bootstraps. (b) Histogram of the root-mean-square error (RMSE) for age predictors trained on 100 random samples pooled from AltumAge’s DNA methylation dataset with the same number of samples as each histone mark. A colored, vertical line shows where in the RMSE distribution the age predictor trained with the histone mark data would lie. (c) Scatter plot of the RMSE of each histone mark age predictor against sample size with a regression line with its 95% confidence interval based on 1000 bootstraps. (d) Three-dimensional scatter plots for samples that were done in triplicates. A dotted black line representing x=y=z is shown alongside a colored, solid regression line with its 95% confidence interval based on 1000 bootstraps. (e) Bar plot showing the intraclass correlation coefficient with error bars representing 95% confidence interval for each histone mark. (f) Point plot with 95% confidence interval based on 1000 bootstraps of the mean absolute error of each histone mark age predictor under added artificial Gaussian noise.

It seems that the sample size is a significant determinant of the performance (Figure 3c), as it is highly negatively correlated with RMSE (r = -0.74, p = 0.059). It is possible that a larger sample size would make histone modification age predictors match or even exceed the performance of DNA methylation age predictors. Therefore, we downloaded all human tissue ChIP-seq samples imputed with Avocado [34]. The original 1814 samples plus 1379 imputed samples were added to 3193 samples. We reran the nested cross-validation adding the imputed samples to the training sets. It has been suggested that the imputed signals contain enough biological information to be useful in several downstream analyses [35]. However, the performance was overall very similar for our age prediction tasks (Supplementary Figure 3g-l). This might suggest that imputed samples are unlikely to help our age predictors; perhaps the performance might be already saturated or the age-related changes are too subtle for Avocado to reconstruct.

In addition to testing the performance on data using features based on the average signal value over gene bodies, we also explored binning the ChIP-Seq data into (1) solely intergenic regions, (2) genes and intergenic regions (whole genome), (3) 20318 CpG dinucleotides common to the Illumina Methylation arrays 27k, 450k, and EPIC, and (4) Horvath’s 353 CpG sites from his pan-tissue DNA methylation age predictor [13]. Despite the different lengths of the genomic loci, that should have not biased our data transformation since we average the signal over the entire bin. Given that heterochromatin is present mainly in noncoding genomic regions, we expected a better performance for the histone modification age predictors based on repressive histone modifications. This is observed — albeit with a minor improvement — for the H3K9me3 age predictor (r = 0.78 vs. 0.74, MAE = 8.67 vs. 8.20, RMSE = 14.52 vs. 15.44) and the H3K36me3 age predictor (r = 0.84 vs. 0.80, MAE = 7.17 vs. 7.86, RMSE = 12.32 vs. 13.48) (Supplementary Figure 3a). The performance is virtually identical for the whole genome setting (Supplementary Figure 3a). The performance of the 20138 CpG sites is similar but slightly worse (Supplementary Figure 3c). Interestingly, it is particularly so for the age predictors of the repressive marks H3K9me3 (r = 0.66 vs. 0.74, MAE = 11.02 vs. 8.20, RMSE = 17.06 vs. 15.44), H3K27me3 (r = 0.90 vs. 0.92, MAE = 7.49 vs. 6.20, RMSE = 12.22 vs. 11.10), and H3K36me3 (r = 0.69 vs. 0.80, MAE = 9.48 vs. 7.86, RMSE = 16.46 vs. 13.48). A similar trend is observed for the age predictors using the histone modification signal from Horvath’s 353 CpG sites, though with overall poorer performance (Supplementary Figure 3d). The methylation status of these CpG sites may interfere with how much epigenetic information can be gained from that particular locus — for instance, if there is high histone acetylation, the gene is almost certainly active. Still, lack of histone mark repression does not mean the gene is active as it might be methylated (though CpG methylation and histone repression typically are well correlated). By binning at different places in the genome, we can see that age-related information is degenerate throughout the genome, as age predictors using inputs from completely different loci perform well. Nonetheless, there are specificities for the performance of histone mark age predictors given the function of the modifications, i.e., some marks perform slightly better or worse than others depending on the loci of the bins.

While it is impossible to directly compare the performance of DNA methylation epigenetic age predictor versus histone mark age predictors without paired data from the same sample, we attempted to make a rough comparison. For such, we ran 100 simulations by randomly drawing the same number of samples of each histone mark from the pan-tissue DNA methylation data set used to create AltumAge [14]. We subjected this random pool to the same nested cross-validation setup using the same machine-learning approach. In the end, we had performance metrics for 100 simulations of each sample size for DNA methylation age predictors. With these results, we compared how well each histone mark age predictor fell into the distribution of DNA methylation age predictors (Figure 3b). If a histone mark age predictor performs well, say over 90th percentile, it means that the reported metric was better than 90 out of the 100 simulations of a DNA methylation age predictor with the same number of samples. Overall, while the H3K9me3 and H3K36me3 age predictors are in the 0th percentile for RMSE, the H3K4me3, H3K9ac, and H3K27ac ones are in the 93rd, 92nd, and 67th percentile, respectively. Similar results were found for Pearson’s correlation and MAE (Supplementary Figure 3e,f). It is worth emphasizing that the AltumAge data was highly skewed towards younger ages, in which DNA methylation age predictors perform better, in contrast to our histone mark data’s more uniform age distribution (Supplementary Figure 2a). Overall, the performance of the histone mark age predictors was approximately in line with the DNA methylation age predictors, with activating histone marks outperforming and repressive histone marks underperforming.

Moreover, we assessed how robust and reliable our histone mark age predictors are. Our data contained samples with biological triplicates, which allowed us to analyze how reliable each histone mark age predictor is to experimental noise. Except for H3K4me1 and H3K36me3, all other histone mark age predictors showed an intraclass correlation coefficient above 0.9 (Figure 3e). In addition to assessing the reliability of the models to experimental variation, we wanted to test how they performed under the addition of artificial noise. For each test fold in the nested cross-validation, we added random Gaussian noise to the test data with up to 1.5 standard deviations in 0.3 standard deviation increments. Even in the most extreme scenario with 1.5 standard deviations, most models’ performance remained similar, except for the H3K4me3 age predictor (Figure 3f). These results show that the age predictors are robust and reliable to experimental and artificial noise.

Lastly, we hypothesized that our choice of the uncertainty-aware ARD regression would give us insights into the epigenetic drift that occurs over time (Supplementary Figure 3g). We expected to see an increased model uncertainty with the sample’s age. Though relatively weak correlations, we did notice a statistically significant relationship for the following age predictors: H3K36me3 (r = 0.14, p = 0.025), H3K4me3 (r = 0.11, p = 0.033), and H3K9me3 (r = 0.14, p = 0.029). Despite the weak correlations, these findings show that the age predictors might be learning the well-described phenomenon of epigenetic drift.

### Inference of pan-tissue histone mark age predictors

While DNA methylation age predictors have been the most used tools to measure age, the insights gained from them into what constitutes epigenetic aging are limited. The most important genes based on the location of the relevant CpG sites are often difficult to relate to the rest of the aging literature. Therefore, we sought to carefully analyze the genes that comprise our histone mark age predictors, i.e., the genes selected after the first step with ElasticNet. In the previous nested cross-validation setup, we trained 10 models in total. However, to be able to interpret the findings more clearly, we ran a single 10-fold cross-validation to choose the best hyperparameter *λ* and trained a single histone modification age predictor with the entirety of the data for each histone.

First, we began by visualizing an upset plot with all histone mark age predictors except for H3K9ac, given its low sample size and poor performance (Figure 4a). The models selected a subset from 341 genes for H3K9me3 up to 1275 for H3K27ac. As expected, the selected genes were often shared across similar histone marks. For instance, the two marks with the most genes in common were H3K27ac and H3K4me3, and the three marks with the most genes in common were H3K27ac, H3K4me3, and H3K4me1. Surprisingly, though few, some selected genes were common across both activating and repressive histone mark age predictors. For the principal components derived from the age predictor genes, the ARD regression decreased the coefficients of only a small subset of genes (about 5%) to zero (Supplementary Figure 4b). This indicates that only a tiny fraction of the 62241 features is sufficient to predict age.

**Figure 4:**
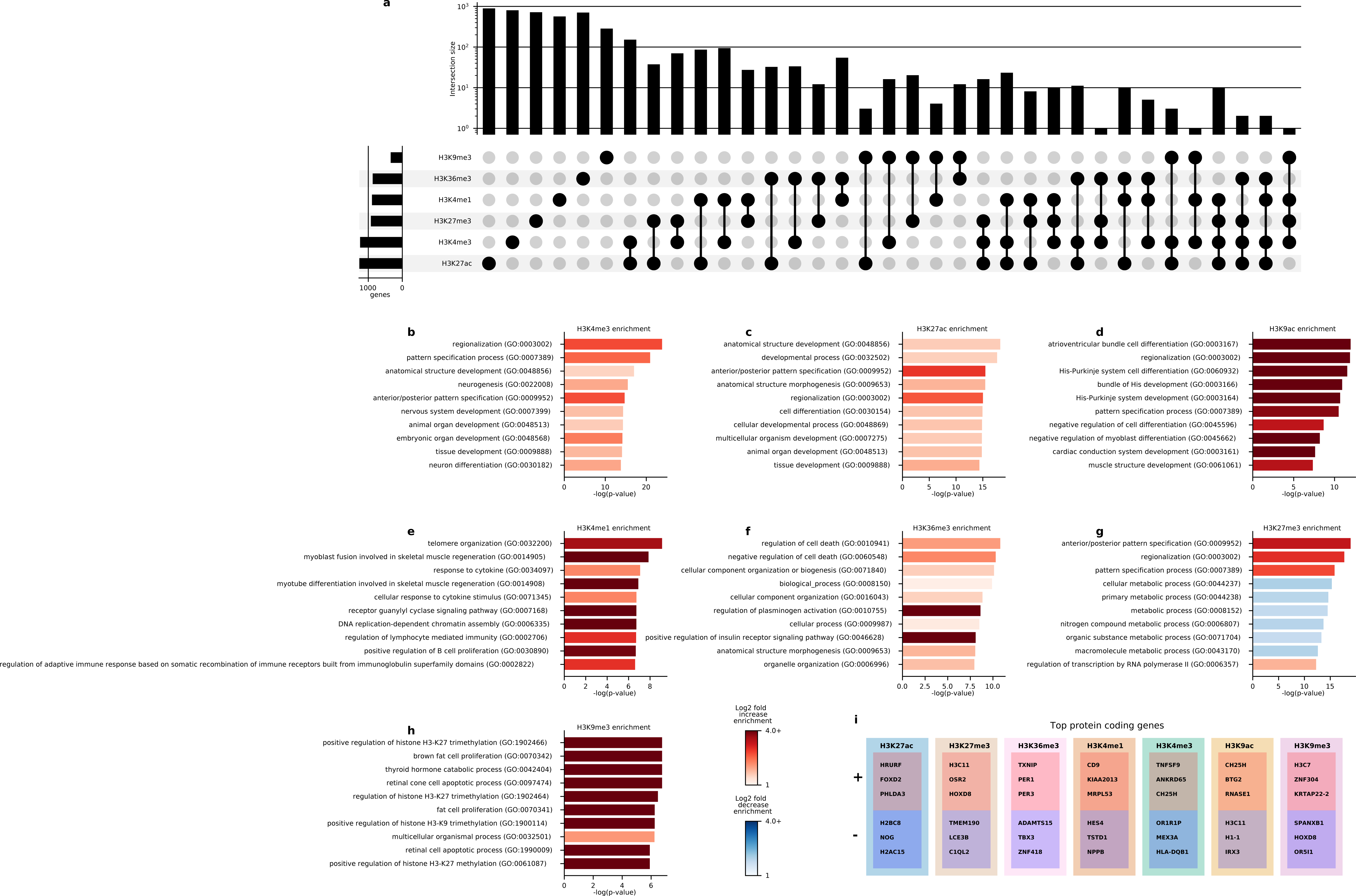
Inference of pan-tissue histone mark age predictors. (a) Upset plot for the subset of genes selected for six of the seven histone modification age predictors. Top 10 gene ontology biological processes from gene set enrichment analysis from Panther DB of H3K4me3 (b), H3K27ac (c), H3K9ac (d), H3K4me1 (e), H3K36me3 (f), H3K27me3 (g), and H3K9me3 (h). (i) Top 3 protein-coding genes for each histone mark age predictor with positive and negative coefficients.

Next, we investigated which pathways were important for the histone mark age prediction. Gene set enrichment analysis (GSEA) can reveal important gene ontology processes which are either over or underrepresented in a set of genes. Using Panther DB [36], we ran seven GSEAs, one for each set of selected genes from the histone mark age predictors (Figure 4b-h). Several developmental pathways are overrepresented in the top 10 GO biological processes. Some examples are regionalization (H3K4me3, H3K27ac, H3K9ac, H3K27me3), pattern specification process (H3K4me3, H3K9ac, H3K27me3), anterior/posterior pattern specification (H3K4me3, H3K27ac, H3K27me3), anatomical structure morphogenesis (H3K27ac, H3K36me3). This brings further evidence to the fact that aging is simply a maladaptive continuation of development, which are fundamental for fitness early in life but whose continuation result in organismal decay. For H3K4me1, processes related to telomere organization, muscle regeneration, and immune response are heavily overrepresented. For H3K9me3, processes related to H3K27me3 regulation and fat proliferation are the most important.

Complementary to the GSEA, we also looked into which ENSEMBL gene biotypes were over or underrep-resented in each model (Supplementary Figure 4a). For such, we used Fisher’s exact test with a Bonferroni correction for the number of gene biotypes (n = 39). Some general trends emerge, with an overrepresentation of micro RNAs (all histone marks), small nuclear RNAs (H3K9me3, H3K27me3, H3K4me1, H3K36me3, H3K27ac), small nucleolar RNAs (H3K9me3, H3K4me3, H3K4me1, H3K36me3, H3K27ac), and miscellaneous RNAs (H3K9me3, H3K4me1, H3K36me3, and H3K27me3). Micro, small nucleolar, and small nuclear RNAs have been linked to several age-related phenomena [19, 23]. Protein-coding genes are vastly underrepresented in the age predictors for all histone modifications besides H3K9ac.

Following the gene set analysis, we looked into the importance of individual genes. We focused on the top three protein-coding genes with the highest positive and negative contributions toward the final age prediction for each age predictor (Figure 4i). Calculating the individual gene importance is possible by inverse-transforming the coefficients of principal components from the ARD regression back into coefficients for the genes. The overarching theme was the importance of histone-coding genes. Amongst those are H1-1, H2AC15, H2BC8, H3C7, H3C11. For activating histone marks, histone genes had a negative coefficient (more histone enrichment translates to lower predicted age), whereas, for the repressive histone modifications, the opposite was true. These genes represent the components of the nucleosome histones H2A, H2B, and H3, and linker histone H1, all of which have been linked to aging [19, 23, 37]. Other relevant age-related genes which contribute towards our histone mark age predictors are NOG, which plays an important role in early development in all germ layers; HOXD8, important in body patterning; TXNIP, whose inhibition can protect against age-related Alzheimer’s disease in mice and whose upregulation causes oxidative stress [38, 39]; PER1 and PER3, circadian-clock genes that are known to be involved in aging [40, 41]; TBX3, which is highly expressed in embryonic stem cells and facilitates cellular reprogramming [42]; TNFSF9, which skews hematopoiesis during aging [43]; BTG2, which drives senescence [44]; CH25H, whose upregulation contributes to the development of osteoarthritis and inflammation in obesity and diabetes [45, 46]. In contrast to several DNA methylation age predictors, histone mark age predictors are enriched in factors that are known to play a role in aging.

### Creation of a pan-histone-mark pan-tissue age predictor

With the results, we had some indications that age predictors trained with similar histone marks behave alike. UMAP clusters activating and repressive marks together (Figure 2a); there is a remarkable correlation between the performance of the age predictors with the number of training samples (Figure 3c); some genes are used by the same models (Figure 4b), and so are similar pathways (Figure 4c-h); the age predictors are generally enriched in similar gene biotypes (Supplementary Figure 4a). Thus, we set out to test whether a pan-histone-mark age predictor with reasonable performance was viable.

First, we made a grid plot contrasting the distribution of Spearman’s correlation between age and each gene for every two histone marks (Figure 5a). As expected, there is usually a positive correlation between activating histone marks and, similarly, between repressive ones. The opposite is true when comparing an activating to a repressive mark. Nevertheless, looking at the density plots, some genes appear to have similar Spearman’s correlation even when contrasting activating and repressive histone modifications. Second, we sorted the protein-coding genes by the highest positive and negative overall Spearman’s correlation with age (Supplementary Figure 5b). Indeed, several histone marks display the same age-related trends. This is because of the generality of the trend in some genes, despite the mainstream assumption that activating and repressive histone marks change in opposite directions. This further supports the hypothesis that a histone mark age predictor trained on one type of histone modification could plausibly predict age with another histone mark as input.

**Figure 5:**
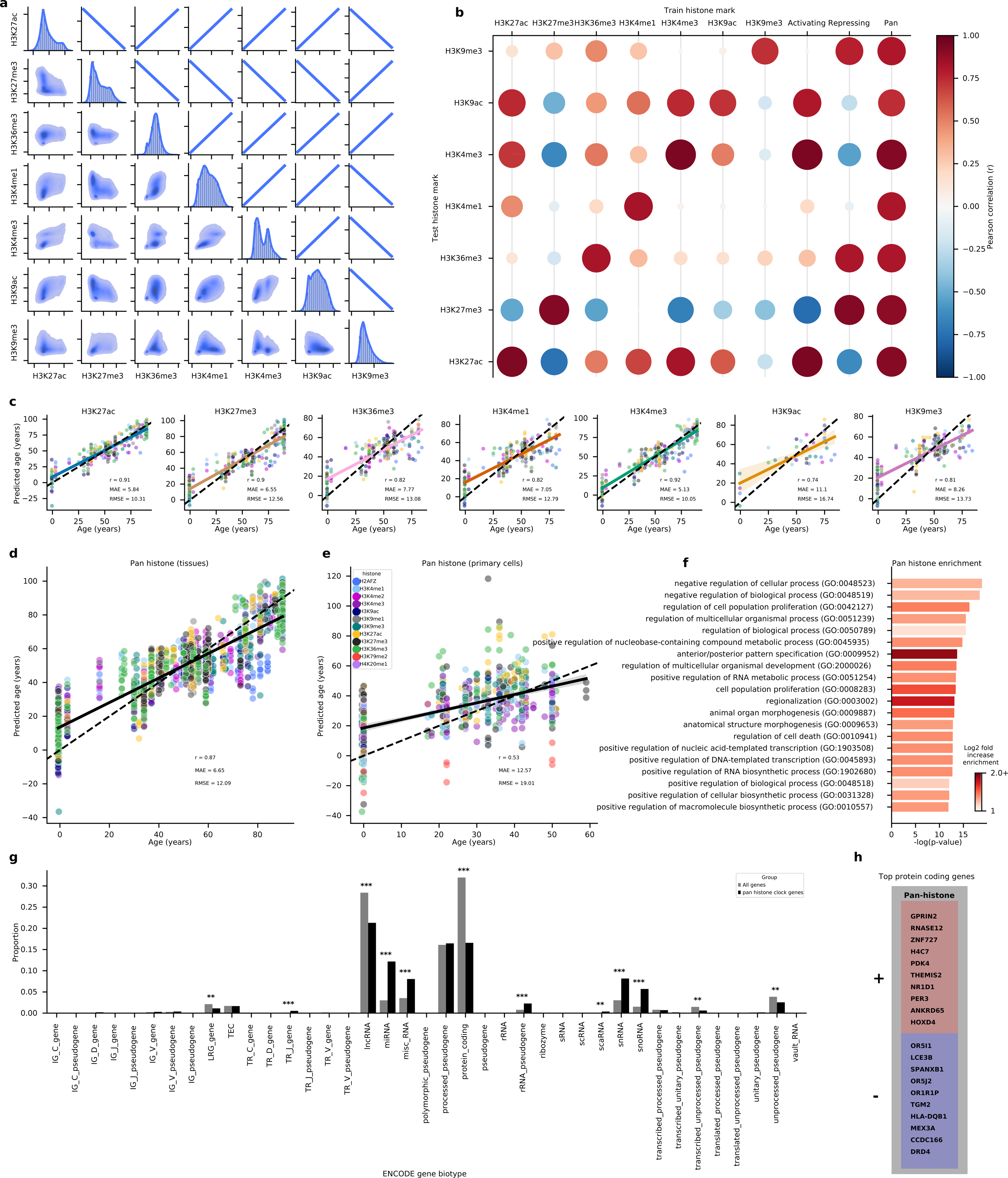
Creation of a pan-histone-mark pan-tissue age predictor. (a) Grid plot with Spearman’s correlation of the histone modification signal for a particular gene over age. Density plots, histograms, and regression lines show the direction of the correlation. (b) Bubble plot of Pearson’s correlation coefficient when age predictors are trained on different histone marks from the ones they aim to predict. Scatter plot of the predicted age of the pan-histone-mark age predictor versus real age using genes as features stratified by histone modification (c) or grouped together (d). Each of the 10 test folds of the nested cross-validation is shown in a different color. A dotted black line representing x=y is shown alongside a colored, solid regression line with its 95% confidence interval based on 1000 bootstraps. (e) Similarly, a scatter plot of the pan-histone-mark age predictor trained using all of the tissue sample data to predict the age of primary cells from 11 different histone marks. Each color represents a distinct histone modification. (f) Top 20 gene ontology biological processes from gene set enrichment analysis from Panther DB for the selected genes from the pan-histone-mark predictor. (g) Bar plot with the proportion of ENSEMBL’s gene biotype for the selected genes in each histone mark age predictor. P-values were rectified with Bonferroni’s correction (*, p *<* 0.01; **, p *<* 0.001; ***, p *<* 0.0001;). (h) Top 10 protein-coding genes for the pan-histone-mark age predictor with positive and negative coefficients.

Next, we reran our nested cross-validation but rather than only predicting the histone mark’s test fold of interest, we instead predicted the test fold of all histone marks. As expected, each histone mark age predictor performs the best when predicting the age of the histone mark with which it was trained (Figure 5b). Interestingly, however, several histone mark age predictors can use other similar histone marks as input with decent performance. For instance, the age predictor trained on H3K27ac performs well (r *>* 0.75) when presented with H3K4me3 and H3K9ac data. The age predictor trained on H3K4me3 performs similarly well when presented with H3K27ac and H3K9ac data. Conversely, the age predictor trained on H3K27me3 has highly negative correlations when predicting the age with the three activating marks.

To test whether the models simply capture “on” or “off” information of a gene rather than a histone-specific signature, we reran the nested cross-validation, flipping the input sign to the ARD regressor. If that was the case, then an age predictor trained on an activating histone mark could theoretically predict the age for a repressive histone mark with the negative of the input signal and vice-versa. This is true despite less significant correlation values across the board (Supplementary Figure 5a). For instance, the flipped H3K4me3 age predictor can predict on H3K27me3 data (r *>* 0.6), and the flipped H3K27me3 age predictor can predict on H3K27ac (r *>* 0.6) and H3K4me3 (r *>* 0.35). This shows that the ChIP-seq signal can be viewed as a sliding scale from repression to activation for some age predictors. This led us to rerun the nested cross-validation, using for training either all activating or all repressive marks plus H3K36me3 — as it is also associated with heterochromatin [47]. Indeed, activating and repressive histone age predictors perform exceedingly well (r *>* 0.8) on activating and repressive histone marks, respectively.

However, one of the most striking observations is that some histone mark age predictors — H3K4me1 and H3K9ac — have positive correlations for six of the seven histone marks. This led us to believe that while some genes contribute towards age prediction based on information akin to a sliding scale of activation and repression, others must supply information differently. Likely, it would be based on age-related cross-histone epigenetic information, which would make creating a pan-histone pan-tissue histone modification age predictor viable. Therefore, we reran the nested cross-validation, using all seven types of histone mark for training or testing, again separating the folds by biosample. The performance for all seven histone marks roughly matches the one from the histone mark-specific age predictor, if not slightly better (Figure 5c). The pan-histone age predictor has a Pearson’s r of 0.87, MAE of 6.65 years, and RMSE of 12.09 years). To further test the generalizability of our pan-histone age predictor, we tested it in untouched data thus far from 568 primary cells spanning 12 histone marks taken from the ENCODE project (Figure 5d). The performance of an age predictor trained on tissues is expected to drop given that the *in vitro* milieu and passaging can induce changes akin to aging [48]. The model’s performance on these cultured cells is still significant (r = 0.53, MAE = 12.57, RMSE = 19.01), as the performance of well-known DNA methylation age predictors is modified by several *in vitro* variables [49].

Next, we looked into some critical pathways and genes for the pan-histone age predictor. Moreover, using Panther DB [36], we analyzed the gene ontology biological processes which were either over or underrepresented in the set of genes selected with the ElasticNet step (Figure 5f). Most gene ontology terms are related to developmental and transcriptional processes. Like histone-specific age predictors, the gene set is underrepresented in protein-coding genes and overrepresented in micro, miscellaneous, small nuclear, and small nucleolar RNA (p-adjusted *<* 0.0001, Figure 5g). Similar genes appear in the histone-specific age predictors when looking at the protein-coding genes with the highest positive and negative contribution to the pan-histone age predictor (Figure 5h). Amongst those are H4C7, a component of nucleosomes; HOXD4, important in body patterning; NR1D1 and PER3, involved in regulating circadian rhythms. Curiously, several olfactory receptor genes also appeared important for age prediction. Recently, this sense has been implicated in lifespan regulation in worms [50].

## Discussion

The allure of DNA methylation age predictors, given their remarkable precision, has significantly redirected the focus in aging research towards epigenetics. Previously, the foundation for epigenetic age predictors predominantly rested on DNA methylation. However, recent advancements have led to the inception of a model grounded in chromatin accessibility, which has shown promising results [51]. Until this study, a predictor based on histone marks was a void in the research domain. Our work elucidates that ChIP-Seq data, derived from seven histone marks, can underpin the formulation of highly precise age predictors. A simulated analysis indicates that histone mark age predictors, given comparable sample sizes, could potentially surpass their DNA methylation counterparts in terms of accuracy, especially concerning activating histone marks. Fundamentally, our findings champion the age-associated dynamics of histone modifications as potent aging biomarkers, integral for devising resilient age predictors.

While DNA methylation age predictors are commendable in performance, their interpretability often remains obscure. Crucial CpG sites integral to these models often pertain to genes of elusive function or ones whose aging impact is dubious. For instance, a recent endeavor to construct a lifespan DNA methylation predictor highlights a CpG site near BCL11B as vital [52]. However, its attenuation scarcely affects the epigenetic age in mice. Another CpG site related to the ELOV2 gene, crucial in fatty acid metabolism, while having potential health implications, bears a tenuous link with aging. In contrast, our histone mark age predictors teem with well-established aging biology. These models underscore pivotal genes associated with development, inflammation, senescence, and stem cell sustenance, among others. Interestingly, some pathways enriched in our predictors mirror those in a recent accelerated aging mouse model [53].

Moreover, our results pave the way for formulating intriguing longevity interventions. Although the significance of genes in our models is inherently correlative, it would be fascinating to explore interventions targeting these gene modifications. For example, yeast lifespan has been shown to extend upon overexpressing certain histone proteins [54]. Additionally, medications influencing histone modifications have demonstrated their potential in governing health and lifespan in specific organisms. Examples include NAD+ precursor supplementation impacting sirtuin enzymes [55, 56] and the effects of alpha-ketoglutarate on certain histone demethylases [57–59]. Recent revelations suggest that compounds influencing histone modifications can rejuvenate cells *in vitro* [60, 61]. Thus, our models’ top hits and interventions affecting histone modifications might chart innovative paths in aging research.

One of this paper’s pivotal contributions is the demonstration that a unified age predictor can adeptly decipher age from various epigenetic data types. Merging different data modalities for a unified model has precedent [51]. Yet, our results suggest distinct histone marks encapsulate consistent age-related data. A case in point is the effectiveness of a predictor trained on H3K4me3 data when applied to H3K27ac data. We have crafted a pan-histone, pan-tissue age predictor compatible across numerous histone marks both in tissue and *in vitro* primary cells. Remarkably, this predictor can seamlessly assimilate multiple data types without necessitating model weight adjustments, underlining the significance of age-related patterns over specific genomic locus changes.

## Conclusion

In summary, this study presents a holistic inspection of age-related changes in histone marks. We introduce innovative age predictors, underscoring the potential to utilize a solitary model capable of handling varied data modalities for age prediction. A limitation to consider is the dearth of quality ChIP-Seq data beyond the ENCODE project. High-quality ChIP-Seq data is scarce compared to DNA methylation and is needed for the appropriate processing of the bigWig files necessary for our models. Optimally, our predictors would be cross-validated using samples adhering to ENCODE guidelines. Nonetheless, the prohibitive cost of ChIP-Seq, compared to the prevalent methylation arrays, has constrained the creation of expansive datasets at this study’s juncture. We are optimistic about future experiments building upon the insights this paper offers.

## Methods

### Data

To generate and interpret the age predictors, we collected 1814 human tissue ChIP-Seq samples from the ENCODE project in the bigWig format [21, 22] for the histone modifications H3K4me3 (n = 359), H3K27ac (n = 359), H3K27me3 (n = 291), H3K4me1 (n = 264), H3K36me3 (n = 257), H3K9me3 (n = 248), and H3K9ac (n = 36). The data was generated from Bradley Bernstein’s lab at the Broad Institute, John Stamatoyannopoulos’s lab at UW, Joseph Costello’s lab at UCSF, and Bing Reng’s lab at UCSD.

We generated a feature matrix by averaging the negative log10 of p-values signal across all nucleotides per gene body according to the Homo Sapiens Ensembl annotation 105 [26]. Of note, the p-values are already -log10-transformed in the ENCODE bigWigs. Then, these averages were arcsinh-transformed. The summary of the transformation is as follows:

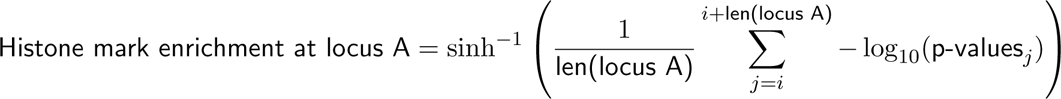

Samples in which more than 10% of the features were unavailable were discarded. Then, missing values for features were encoded as 0. In Section 2 specifically, we also tested averaging the negative log of p-values signal across intergenic regions, genes, and intergenic regions (whole genome), 20318 CpG dinucleotides common to the Illumina Methylation arrays 27k, 450k, and EPIC, and Horvath’s 353 CpG sites from his pan-tissue DNA methylation age predictor [13].

Embryonic samples had their age encoded as 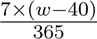, where *w* is the gestational week, and 40 is the number of weeks of a normal gestation [62]. Therefore, some samples had negative ages. Some samples whose age was described as 90-plus by the ENCODE project, likely for anonymity reasons, were encoded as 90 years.

The distribution of the data is represented in Supplementary Figure 2.

Moreover, to test the performance *in vitro* of our pan-histone pan-tissue age predictor, we gathered another 568 samples of primary cells spanning 12 histone marks (H2AFZ, H3K4me1, H3K4me2, H3K4me3, H3K9ac, H3K9me1, H3K9me3, H3K27ac, H3K27me3, H3K36me3, H3K79me2, H4K20me1) taken from the ENCODE project [21, 22].

Lastly, to assess the tentative increase in performance that could arise from increasing the sample size through imputation, we further downloaded all available Avocado-imputed samples from ENCODE for the seven prominent histone marks we analyzed in the paper, 197 of each, totaling 1379 samples.

### Age predictor performance evaluation

For all experiments assessing the performance of the age predictors, the setup consisted of 10-fold nested cross-validation with observations from the same biosample remaining in the same folds to not artificially boost the performance. Nine folds are used for internal 9-fold cross-validation to select the appropriate hyperparameter. The nine folds are used for training the model, which is tested in the remaining external fold. Cancer samples were always removed.

Our age predictors consist of three steps. The feature selection method employs an ElasticNet model with a 0.9 L1 to L2 proportion to choose *p* features whose absolute coefficient is above zero. The only hyperparameter *λ* in the age predictors is the strength of regularization of the feature-selection ElasticNet, which controls how many variables are chosen. The tested hyperparameter values were *λ ∈ {*0.1, 0.05, 0.01, 0.005, 0.001*}*. Secondly, to improve the robustness of the age predictor, we applied PCA calculated with a truncated support vector decomposition to generate (*p^′^ −* 1) principal components. Finally, we used an automatic relevance determination regression [63], a form of regularized Bayesian regression, to predict age. All of the steps above were created with sklearn library in Python. If not mentioned otherwise, all other hyperparameters were the standard ones in the package.

To roughly compare the performance of using histone mark data to DNA methylation data [14], we used AltumAge’s pan-tissue dataset and pooled 100 random samples of the same size as the sample size for each histone modification. Then, we ran the same three steps we used to create our histone modification age predictors and ended up with 100 values for each performance metric. With this information, we compared where the performance of the histone mark age predictors fell within the distribution for the DNA methylation age predictors taking sample size into account. However, it must be emphasized that the ENCODE ChIP-seq dataset does not have the same age and tissue distribution as AltumAge’s DNA methylation dataset.

In the setup to assess the performance of our models with the addition of the Avocado-imputed samples, whenever a model was trained on a particular histone mark, the imputed samples were added to the training set (with tumors removed) for both each inner fold of the nested cross-validation and when training the model to predict each test set of the cross-validation. The imputed samples were only added to try to improve the performance of predicting the age of actual samples rather than attempting to predict the age of imputed samples.

### Age predictor interpretation

To interpret a particular age predictor, we ran a 10-fold cross-validation to select the best regularization hyper-parameter in the ElasticNet feature selection step. Then, we trained the age predictor using the entirety of the data in each setting. For the gene set enrichment analysis, we selected the genes that passed feature selection: H3K4me3 (*p^′^* = 1240), H3K27ac (*p^′^* = 1275), H3K27me3 (*p^′^* = 922), H3K4me1 (*p^′^* = 892), H3K36me3 (*p^′^* = 870), H3K9me3 (*p^′^* = 341), H3K9ac (*p^′^* = 102), and pan-histone (*p^′^* = 3739). To determine the individual contribution of each feature towards the final age prediction, we inverse-transformed the coefficients of principal components from the automatic relevance determination regression back into coefficients for the genes.

### Statistical Analysis

The p-values associated with Pearson’s and Spearman’s correlation coefficients were obtained using the functions pearsonr and spearmanr from the python package scipy.

To create the UMAP and PCA plots in Figure 2 and Supplementary Figure 2, we ran Python’s dynamo package function dyn.tl.reduceDimension with either UMAP or PCA as the basis with standard parameters.

The intraclass correlation coefficient measures how well multiple measurements agree with one another and is used to assess model reliability. To calculate the intraclass correlation coefficient, we used Python’s pingouin package with a single-rater, absolute-agreement, two-way random-effects model per [64] guidelines and similarly to [31].

Notebooks with the analyses and a complete list of the python package versions are also available on our GitHub (URLXXXXX).

## Declarations

### Ethics approval and consent to participate

Not applicable.

### Consent for publication

Not applicable.

### Availability of data and materials

After publication, the code to rerun our results will be available on our GitHub (URLXXXXX). It takes about three weeks for all scripts to run on an Amazon AWS ml.t3.2xlarge instance.

The data will be available after the publication on Zenodo, for review we have provided the datasets on Google Drive (URLXXXXX).

### Author Contributions

L.P.D.L.C. conceived of the presented idea, devised the methodology, ran experiments, and wrote the first draft of the manuscript. M.H.A. ran experiments and collected the datasets. S.H. assisted with the analysis and biological interpretation of the results and contributed to the final manuscript. E.L. assisted with the analysis and biological interpretation of the results and contributed to the final manuscript. R.S. supervised the project, devised the methodology, and contributed to the final manuscript.

## Acknowledgements

The title of this paper was inspired by [13].

---

**Supplementary Figure 1:**
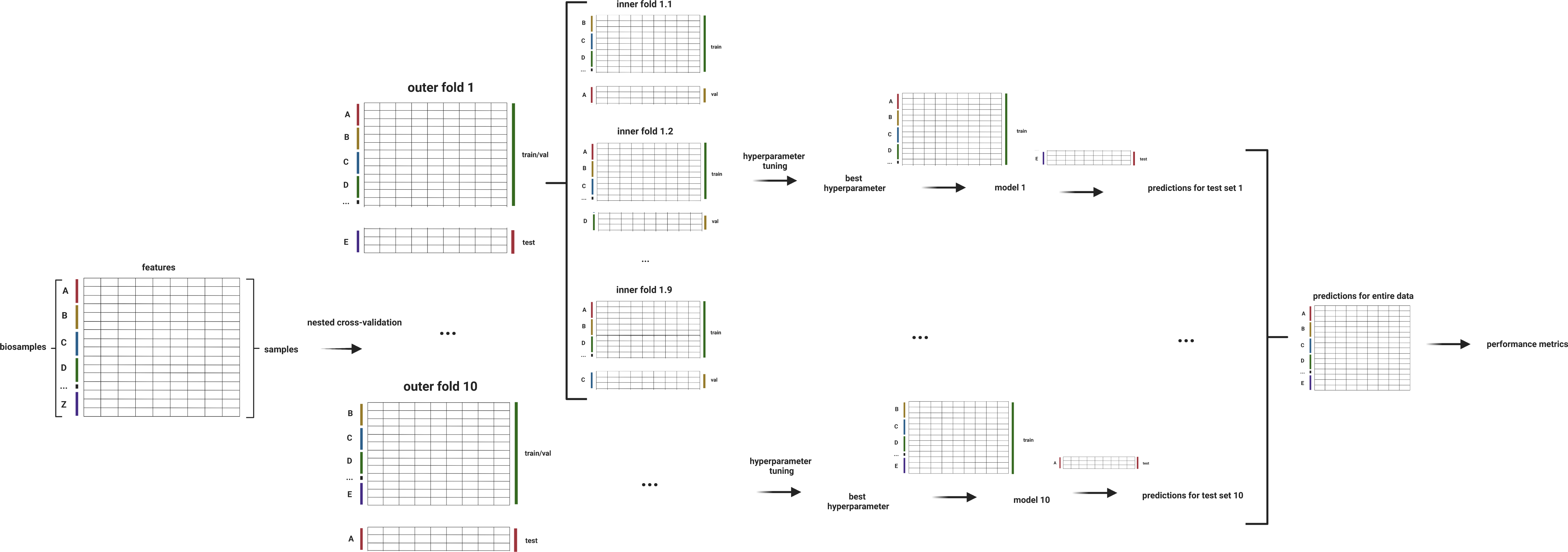
Setup for the nested cross-validation of the different histone mark age predictors. First, the data is split into ten folds. Each fold is divided into training and validation (9/10 of the data) and test (1/10 of the data), always maintaining observations from the same biosample. There is another 9-fold cross-validation using the training and validation part of the data for hyperparameter tuning. When the best hyperparameter is found, the age predictor is trained on the entire training and validation data and used to predict the remaining test set. After this is done for all ten folds, there is a prediction for each observation in the entire data set. Therefore, performance metrics can be calculated. Image created with BioRender.

**Supplementary Figure 2:**
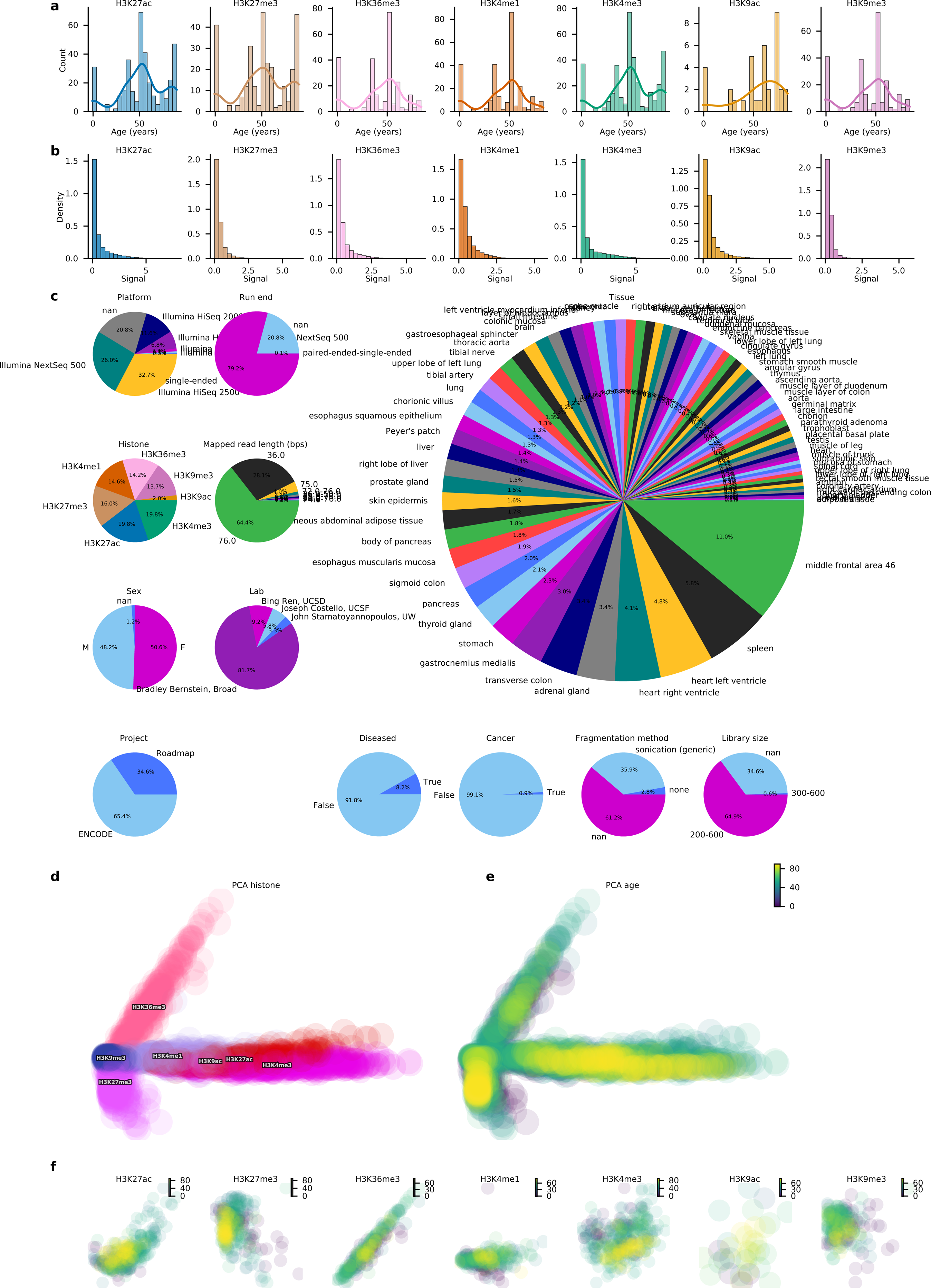
(a) Histogram of the number of samples per histone modification over age. (b) Histogram of the density of transformed signal values per gene. (c) Pie plots with the distribution of the 1814 ChIP-Seq samples by platform, run end, tissue, histone, mapped read length, sex, lab, project, disease status, cancer status, fragmentation method, and library size. (d, e) Principal component analysis (PCA) plot of the dataset grouped by histone modification (d) and age (e) showing the first and second principal components (9.3% and 6.1% of the explained variance, respectively). (f) PCA colored by age for each histone modification.

**Supplementary Figure 3:**
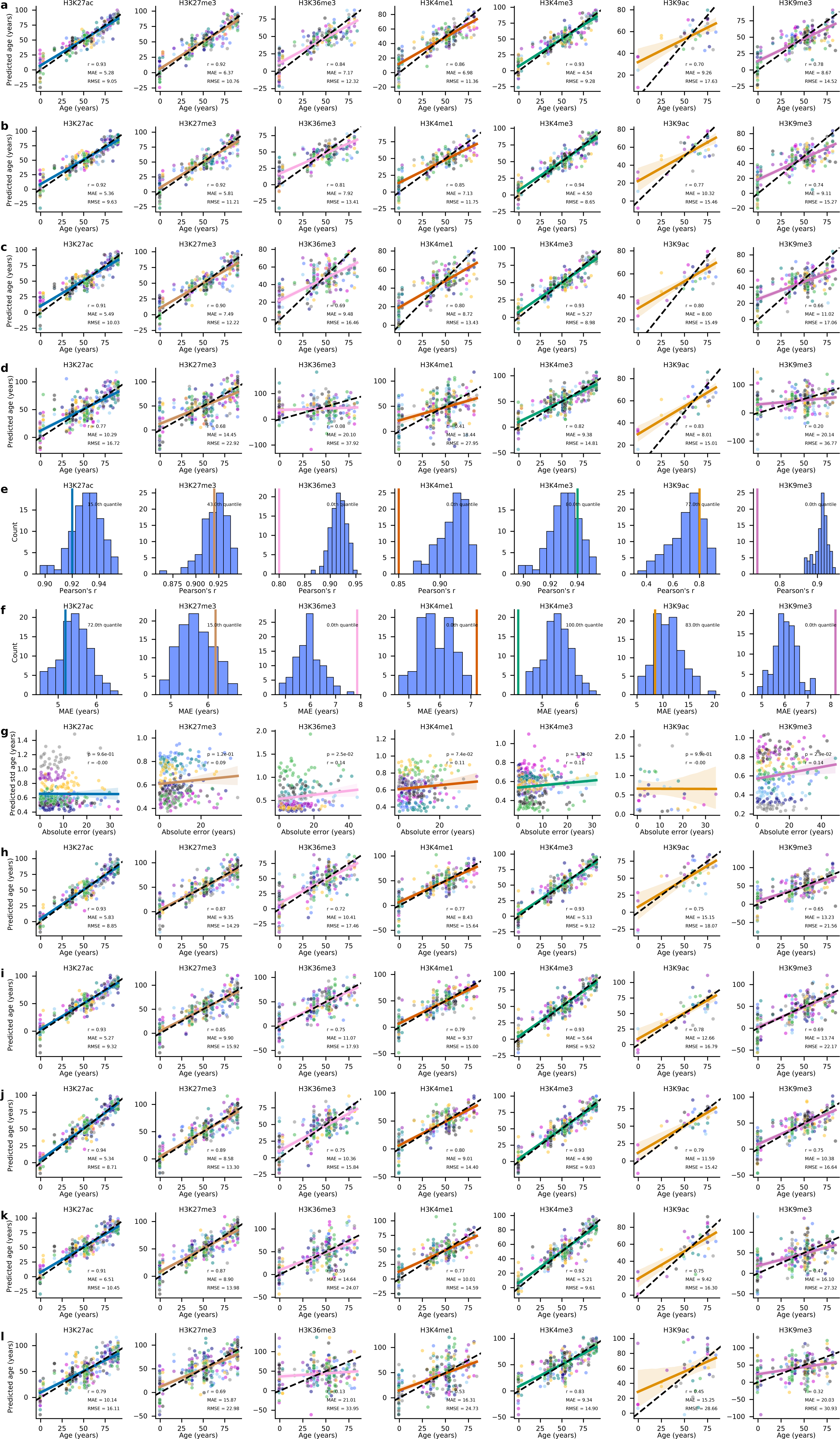
Scatter plot of the predicted age versus real age of each histone mark age predictor using (a) intergenic regions, (b) genes and intergenic regions (whole genome), (c) 20318 CpG dinucleotides common to the Illumina Methylation arrays 27k, 450k, and EPIC, and (d) Horvath’s 353 CpG sites from his pan-tissue DNA methylation age predictor [13] as features. Each of the 10 test folds of the nested cross-validation is shown in a different color. A dotted black line representing x=y is shown alongside a colored, solid regression line with its 95% confidence interval based on 1000 bootstraps. Histogram of Pearson’s correlation (e) and median absolute error (MAE) (f) for age predictors trained on 100 random samples pooled from AltumAge’s DNA methylation dataset with the same number of samples as of each histone mark. A colored, vertical line shows where in the RMSE distribution the age predictor trained with the histone mark data would lie. (g) Scatter plot of the predicted standard deviation versus real age of each histone mark age predictor using genes as features. Each of the 10 test folds of the nested cross-validation is shown in a different color. A colored, solid regression line with its 95% confidence interval based on 1000 bootstraps. Scatter plot of the predicted age versus real age of each histone mark age predictor trained in addition with Avocado-imputed samples [34] using (h) gene bodies, (i) intergenic regions, (j) genes and intergenic regions (whole genome), (k) 20318 CpG dinucleotides common to the Illumina Methylation arrays 27k, 450k, and EPIC, and (l) Horvath’s 353 CpG sites from his pan-tissue DNA methylation age predictor [13] as features. Each of the 10 test folds of the nested cross-validation is shown in a different color. A dotted black line representing x=y is shown alongside a colored, solid regression line with its 95% confidence interval based on 1000 bootstraps.

**Supplementary Figure 4:**
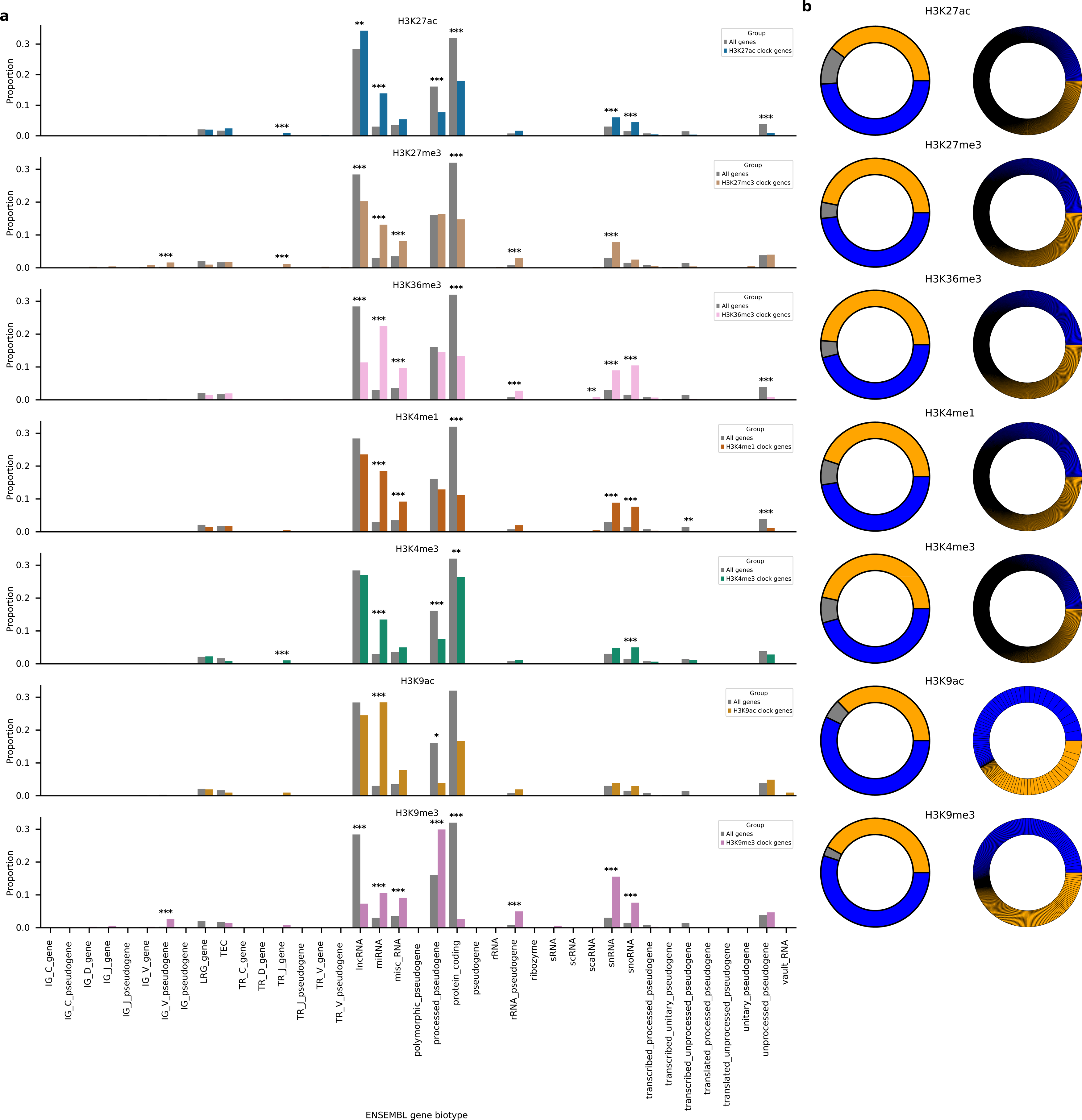
(a) Bar plot with the proportion of ENSEMBL’s gene biotype for the selected genes in each histone mark age predictor. P-values were rectified with Bonferroni’s correction (*, p *<* 0.01; **, p *<* 0.001; ***, p *<* 0.0001;). (b) Doughnut plots for each histone modification age predictor. On the left, the proportion of principal components whose coefficients were positive (yellow), zero (gray), or negative (blue) is displayed; on the right, the weight of each gene to the total prediction is shown, with positive genes with positive coefficient in yellow and negative in blue.

**Supplementary Figure 5:**
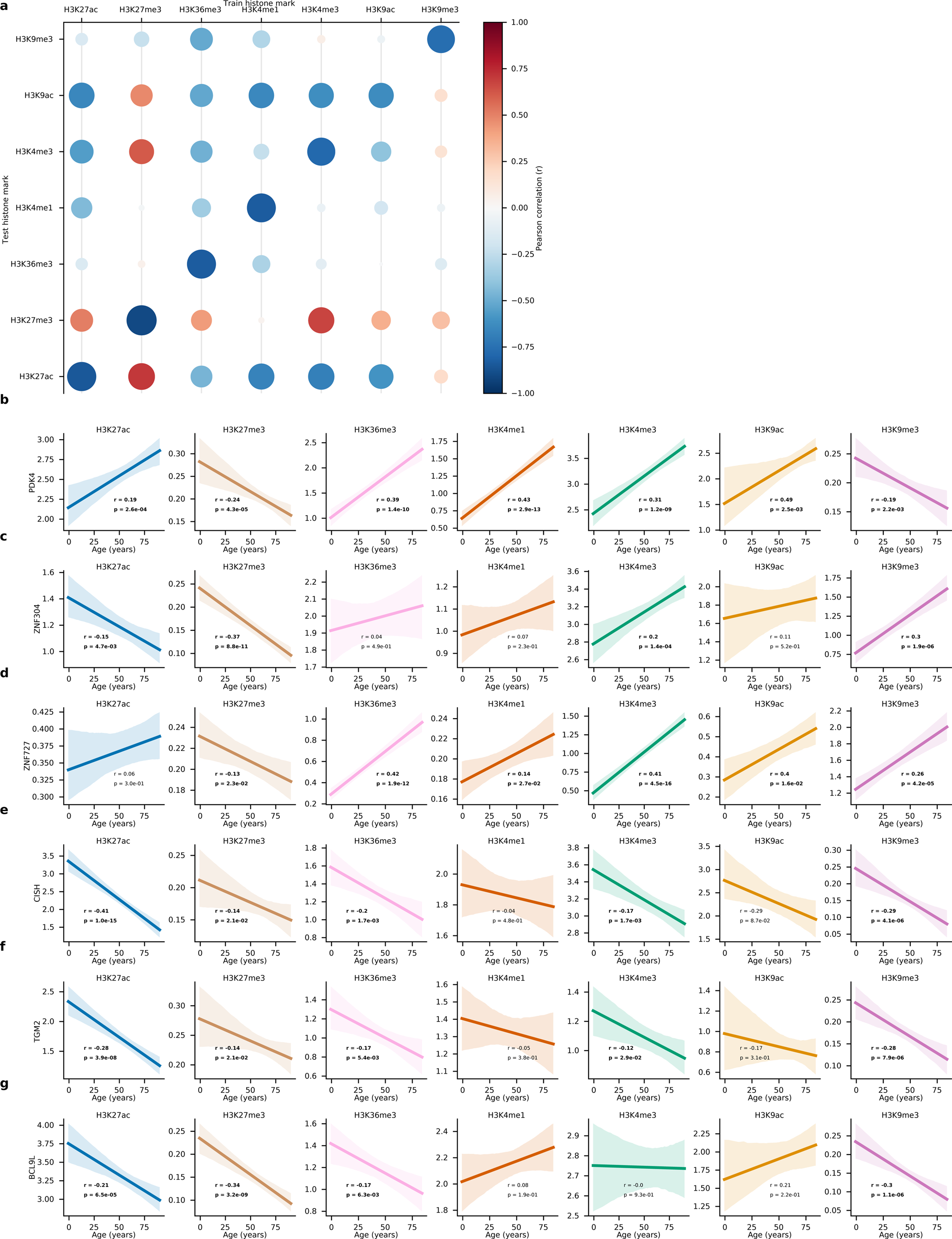
(a) Bubble plot of Pearson’s correlation coefficient when age predictors are trained on certain histone marks and attempt to predict others but with the negative value of the input to the ARD regression part of the age predictor model. (b-g) Regression plots of age versus histone mark signal values for six genes generally go up or down with age. Shaded is the 95% regression confidence interval based on 1000 bootstraps.

## References

[1] Carlos López-Otın, et al. “The hallmarks of aging”. In: Cell 153.6 (2013), pp. 1194–1217.

[2] Carlos Ĺopez-Otın, et al. “Hallmarks of aging: An expanding universe”. In: Cell (2023).

[3] Ruiye Chen, et al. “Biomarkers of ageing: Current state-of-art, challenges, and opportunities”. In: MedComm– Future Medicine 2.2 (2023), e50.

[4] Luigi Ferrucci, et al. “Measuring biological aging in humans: A quest”. In: Aging cell 19.2 (2020), e13080.

[5] Fedor Galkin, et al. “Biohorology and biomarkers of aging: Current state-of-the-art, challenges and opportunities”. In: Ageing Research Reviews 60 (2020), p. 101050.

[6] Evgeny Putin, et al. “Deep biomarkers of human aging: application of deep neural networks to biomarker development”. In: Aging (Albany NY) 8.5 (2016), p. 1021.

[7] Polina Mamoshina, et al. “Population specific biomarkers of human aging: a big data study using South Korean, Canadian, and Eastern European patient populations”. In: The Journals of Gerontology: Series A 73.11 (2018), pp. 1482–1490.

[8] Marjolein J Peters, et al. “The transcriptional landscape of age in human peripheral blood”. In: Nature communications 6.1 (2015), pp. 1–14.

[9] Jason G Fleischer, et al. “Predicting age from the transcriptome of human dermal fibroblasts”. In: Genome biology 19.1 (2018), pp. 1–8.

[10] Nicholas Holzscheck, et al. “Modeling transcriptomic age using knowledge-primed artificial neural networks”. In: npj Aging and Mechanisms of Disease 7.1 (2021), pp. 1–13.

[11] Elisa Ferrari, et al. “A deep neural network provides an ultraprecise multi-tissue transcriptomic clock for the short-lived fish Nothobranchius furzeri and identifies predicitive genes translatable to human aging”. In: bioRxiv (2022), pp. 2022–11.

[12] Dmitrii Kriukov, et al. “Longevity and rejuvenation effects of cell reprogramming are decoupled from loss of somatic identity”. In: bioRxiv (2022), pp. 2022–12.

[13] Steve Horvath. “DNA methylation age of human tissues and cell types”. In: Genome biology 14.10 (2013), pp. 1–20.

[14] Lucas Paulo de Lima Camillo, Louis R Lapierre, and Ritambhara Singh. “A pan-tissue DNA-methylation epigenetic clock based on deep learning”. In: npj Aging 8.1 (2022), pp. 1–15.

[15] Steve Horvath and Kenneth Raj. “DNA methylation-based biomarkers and the epigenetic clock theory of ageing”. In: Nature Reviews Genetics 19.6 (2018), pp. 371–384.

[16] Christopher G Bell, et al. “DNA methylation aging clocks: challenges and recommendations”. In: Genome biology 20 (2019), pp. 1–24.

[17] Alexandre Trapp, Csaba Kerepesi, and Vadim N Gladyshev. “Profiling epigenetic age in single cells”. In: Nature Aging 1.12 (2021), pp. 1189–1201.

[18] Ake T Lu, et al. “Universal DNA methylation age across mammalian tissues”. In: BioRxiv (2021), pp. 2021–01.

[19] Alice E Kane and David A Sinclair. “Epigenetic changes during aging and their reprogramming potential”. In: Critical reviews in biochemistry and molecular biology 54.1 (2019), pp. 61–83.

[20] Na Yang, et al. “A hyper-quiescent chromatin state formed during aging is reversed by regeneration”. In: Molecular Cell 83.10 (2023), pp. 1659–1676.

[21] Encode Project Consortium, et al. “An integrated encyclopedia of DNA elements in the human genome”. In: Nature 489.7414 (2012), p. 57.

[22] Yunhai Luo, et al. “New developments on the Encyclopedia of DNA Elements (ENCODE) data portal”. In: Nucleic acids research 48.D1 (2020), pp. D882–D889.

[23] Lucas Paulo de Lima Camillo and Robert Quinlan. “A ride through the epigenetic landscape: aging reversal by reprogramming”. In: GeroScience 43.2 (2021), pp. 463–485.

[24] Moyra Lawrence, Sylvain Daujat, and Robert Schneider. “Lateral thinking: how histone modifications regulate gene expression”. In: Trends in Genetics 32.1 (2016), pp. 42–56.

[25] Andrew J Bannister and Tony Kouzarides. “Regulation of chromatin by histone modifications”. In: Cell research 21.3 (2011), pp. 381–395.

[26] Fiona Cunningham et al. “Ensembl 2022”. In: Nucleic acids research 50.D1 (2022), pp. D988–D995.

[27] Gregory Hannum, et al. “Genome-wide methylation profiles reveal quantitative views of human aging rates”. In: Molecular cell 49.2 (2013), pp. 359–367.

[28] Andrei E Tarkhov, Kirill A Denisov, and Peter O Fedichev. “Aging clocks, entropy, and the limits of age-reversal”. In: bioRxiv (2022).

[29] Andrei E Tarkhov, et al. “Nature of epigenetic aging from a single-cell perspective”. In: bioRxiv (2022).

[30] Hui Zou and Trevor Hastie. “Regularization and variable selection via the elastic net”. In: Journal of the royal statistical society: series B (statistical methodology) 67.2 (2005), pp. 301–320.

[31] Albert T Higgins-Chen, et al. “A computational solution for bolstering reliability of epigenetic clocks: Implications for clinical trials and longitudinal tracking”. In: Nature aging 2.7 (2022), pp. 644–661.

[32] Fedor Galkin, et al. “DeepMAge: a methylation aging clock developed with deep learning”. In: Aging and disease 12.5 (2021), p. 1252.

[33] Benjamin L Kidder, Gangqing Hu, and Keji Zhao. “ChIP-Seq: technical considerations for obtaining high-quality data”. In: Nature immunology 12.10 (2011), pp. 918–922.

[34] Jacob Schreiber, et al. “Avocado: a multi-scale deep tensor factorization method learns a latent representation of the human epigenome”. In: Genome biology 21.1 (2020), pp. 1–18.

[35] Jacob Schreiber, Jeffrey Bilmes, and William Stafford Noble. “Completing the ENCODE3 compendium yields accurate imputations across a variety of assays and human biosamples”. In: Genome biology 21.1 (2020), pp. 1–13.

[36] Paul D Thomas, et al. “PANTHER: Making genome-scale phylogenetics accessible to all”. In: Protein Science 31.1 (2022), pp. 8–22.

[37] Sangita Pal and Jessica K Tyler. “Epigenetics and aging”. In: Science advances 2.7 (2016), e1600584.

[38] Xue Zhang, et al. “Salidroside ameliorates Parkinson’s disease by inhibiting NLRP3-dependent pyroptosis”. In: Aging (Albany NY) 12.10 (2020), p. 9405.

[39] Rongbin Zhou, et al. “Thioredoxin-interacting protein links oxidative stress to inflammasome activation”. In: Nature immunology 11.2 (2010), pp. 136–140.

[40] Janine L Kwapis, et al. “Epigenetic regulation of the circadian gene Per1 contributes to age-related changes in hippocampal memory”. In: Nature communications 9.1 (2018), pp. 1–14.

[41] Shin Yamazaki, et al. “Effects of aging on central and peripheral mammalian clocks”. In: Proceedings of the National Academy of Sciences 99.16 (2002), pp. 10801–10806.

[42] Jianyong Han, et al. “Tbx3 improves the germ-line competency of induced pluripotent stem cells”. In: Nature 463.7284 (2010), pp. 1096–1100.

[43] Qianqiao Tang, et al. “CD137 ligand reverse signaling skews hematopoiesis towards myelopoiesis during aging”. In: Aging (Albany NY) 5.9 (2013), p. 643.

[44] Kan Wang, et al. “SUV39H2/KMT1B inhibits the cardiomyocyte senescence phenotype by down-regulating BTG2/PC3”. In: Aging (Albany NY) 13.18 (2021), p. 22444.

[45] Wan-Su Choi, et al. “The CH25H–CYP7B1–RORα axis of cholesterol metabolism regulates osteoarthritis”. In: Nature 566.7743 (2019), pp. 254–258.

[46] Lucia Russo, et al. “Cholesterol 25-hydroxylase (CH25H) as a promoter of adipose tissue inflammation in obesity and diabetes”. In: Molecular metabolism 39 (2020), p. 100983.

[47] Sophie Chantalat, et al. “Histone H3 trimethylation at lysine 36 is associated with constitutive and facultative heterochromatin”. In: Genome research 21.9 (2011), pp. 1426–1437.

[48] Gabriel Sturm, et al. “A multi-omics longitudinal aging dataset in primary human fibroblasts with mitochondrial perturbations”. In: Scientific Data 9.1 (2022), p. 751.

[49] Gabriel Sturm, et al. “OxPhos defects cause hypermetabolism and reduce lifespan in cells and in patients with mitochondrial diseases”. In: Communications biology 6.1 (2023), p. 22.

[50] Evandro A De-Souza, Maximillian A Thompson, and Rebecca C Taylor. “Olfactory chemosensation extends lifespan through TGF-β signaling and UPR activation”. In: Nature Aging (2023), pp. 1–10.

[51] Cheyenne Rechsteiner, et al. “Development of a novel aging clock based on chromatin accessibility”. In: bioRxiv (2022).

[52] Amin Haghani, et al. “Divergent age-related methylation patterns in long and short-lived mammals”. In: bioRxiv (2022).

[53] Jae-Hyun Yang, et al. “Loss of epigenetic information as a cause of mammalian aging”. In: Cell 186.2 (2023), pp. 305–326.

[54] Sun-Ju Yi and Kyunghwan Kim. “New insights into the role of histone changes in aging”. In: International Journal of Molecular Sciences 21.21 (2020), p. 8241.

[55] Sarah J Mitchell, et al. “Nicotinamide improves aspects of healthspan, but not lifespan, in mice”. In: Cell metabolism 27.3 (2018), pp. 667–676.

[56] Kathryn F Mills, et al. “Long-term administration of nicotinamide mononucleotide mitigates age-associated physiological decline in mice”. In: Cell metabolism 24.6 (2016), pp. 795–806.

[57] Khoa A Tran, Caleb M Dillingham, and Rupa Sridharan. “The role of α-ketoglutarate–dependent proteins in pluripotency acquisition and maintenance”. In: Journal of Biological Chemistry 294.14 (2019), pp. 5408–5419.

[58] Azar Asadi Shahmirzadi, et al. “Alpha-ketoglutarate, an endogenous metabolite, extends lifespan and compresses morbidity in aging mice”. In: Cell metabolism 32.3 (2020), pp. 447–456.

[59] Oleksandr Demidenko et al. “Rejuvant®, a potential life-extending compound formulation with alpha-ketoglutarate and vitamins, conferred an average 8 year reduction in biological aging, after an average of 7 months of use, in the TruAge DNA methylation test”. In: Aging (albany NY) 13.22 (2021), p. 24485.

[60] Jae-Hyun Yang, et al. “Chemically induced reprogramming to reverse cellular aging.” In: Aging 15 (2023).

[61] Wayne Mitchell, et al. “Multi-omics characterization of partial chemical reprogramming reveals evidence of cell rejuvenation”. In: bioRxiv (2023), pp. 2023–06.

[62] Joyce A Martin et al. “Measuring gestational age in vital statistics data: transitioning to the obstetric estimate.” In: National Vital Statistics Reports: From the Centers for Disease Control and Prevention, National Center for Health Statistics, National Vital Statistics System 64.5 (2015), pp. 1–20.

[63] David JC MacKay, et al. “Bayesian nonlinear modeling for the prediction competition”. In: ASHRAE transactions 100.2 (1994), pp. 1053–1062.

[64] Terry K Koo and Mae Y Li. “A guideline of selecting and reporting intraclass correlation coefficients for reliability research”. In: Journal of chiropractic medicine 15.2 (2016), pp. 155–163.

